# Evolution-based design of chorismate mutase enzymes

**DOI:** 10.1101/2020.04.01.020487

**Authors:** William P. Russ, Matteo Figliuzzi, Christian Stocker, Pierre Barrat-Charlaix, Michael Socolich, Peter Kast, Donald Hilvert, Remi Monasson, Simona Cocco, Martin Weigt, Rama Ranganathan

## Abstract

The rational design of enzymes is an important goal for both fundamental and practical reasons. Here, we describe a design process in which we learn the constraints for specifying proteins purely from evolutionary sequence data, build libraries of synthetic genes, and test them for activity *in vivo* using a quantitative complementation assay. For chorismate mutase, a key enzyme in the biosynthesis of aromatic amino acids, we demonstrate the design of natural-like catalytic function with substantial sequence diversity. Further optimization focuses the generative model towards function in a specific genomic context. The data show that sequence-based statistical models suffice to specify proteins and provide access to an enormous space of synthetic functional sequences. This result provides a foundation for a general process for evolution-based design of artificial proteins.

**One-sentence summary:** An evolution-based, data-driven engineering process can build synthetic functional enzymes.

---

Approaches for protein design typically begin with atomic structures and physical models for forces between atoms, but the dramatic expansion of protein sequence databases and the growth of new computational methods (*1-4*) opens new strategies to this problem. Statistical analyses of homologs comprising a protein family have recently enabled successful prediction of protein structure (*5-8*), protein-protein interactions (*9-13*), mutational effects (*14-18*), and for a family of small protein interaction modules, have led to the successful design of artificial amino acid sequences that fold and function in a manner similar to their natural counterparts (*19, 20*). These findings indicate that sequence-based statistical models may represent a general approach for a purely data-driven strategy for the design of complex proteins that can work *in vivo*, in specific organismal contexts. A first key step is to demonstrate the sufficiency of these models for specifying functional proteins.

An approach for evolution-inspired protein design is shown in Figure 1A-C, based on direct coupling analysis (DCA), a method originally conceived to predict contacts between amino acids in protein three-dimensional structures (*1*). The starting point is a large and diverse multiple sequence alignment (MSA) of a protein family, from which we estimate the observed frequencies 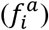 and pairwise co-occurrences 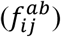 of all amino acids (*a, b*) at positions (*i, j*) – the first- and second-order statistics (Fig. 1A). From these quantities, we infer a model comprising a set of intrinsic amino acid propensities (fields *h*_*i*_(*a*)) and a set of pairwise interactions (couplings *J*_*ij*_ (*a, b*)) that optimally account for the observed statistics (Fig. 1A). This model defines a probability *P* for each amino acid sequence (*a*, ⋯, *a*_*L*_) of length *L*:

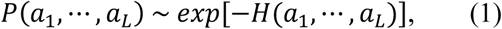

with the Hamiltonian *H*(*a*_1_, ⋯, *a*_*L*_) = − ∑_*i*_ *h*_*i*_(*a*_*i*_) − ∑_*i* < *j*_ *J*_*ij*_(*a*_*i*_, *a*_*j*_) representing a statistical energy (that provides a quantitative log-likelihood score for each sequence, see Methods). Lower energies are associated with higher probability. Monte Carlo (MC) sampling from the model allows for generating non-natural sequence repertoires (Fig. 1B), which can then be screened for desired functional activities (Fig. 1C). If positional conservation and pairwise correlation suffice in general to capture the information content of protein sequences and if the model inference is sufficiently accurate, then the synthetic sequences should recapitulate the functional diversity and properties of natural proteins.

**Figure 1:**
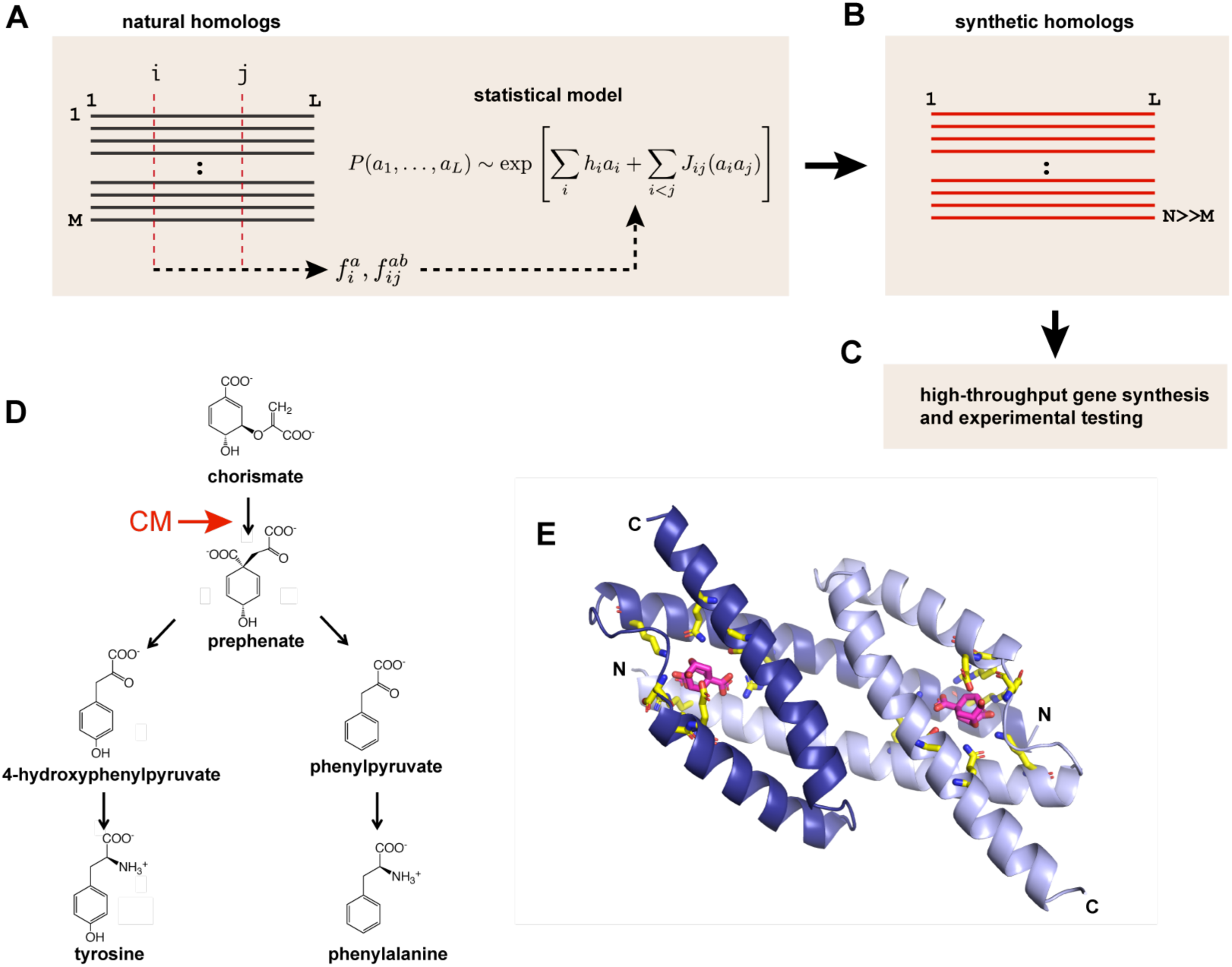
Evolutionary data-driven protein engineering. **A**, a multiple sequence alignment (MSA) of *M* natural homologs provides empirical first and second-order statistics of amino acids 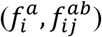, which are used to infer a statistical model with the bmDCA method. The probability of sequence ***a*** = (*a*_1_, …, *a*_*L*_) is an exponential function of a Hamiltonian, or statistical energy, parameterized by intrinsic fields *h*_*i*_(*a*) and couplings *J*_*ij*_(*a, b*) acting on amino acids. **B-C**, the model is used to generate *N* ≫ *M* synthetic sequences that can be tested in a high-throughput assay for desired functions. **D**, chorismate mutase (CM) is an enzyme occurring at the central branch point in the shikimate pathway leading to the synthesis of tyrosine and phenylalanine. **E**, members of the AroQ_α_ and AroQ_δ_ families of CMs fold into a domain-swapped dimer (PDB ID 1ECM). Active site residues are shown with yellow stick bonds and arise from both subunits (dark and light blue). A bound substrate analog is shown in magenta.

To test this process, we chose the AroQ family of chorismate mutases (CMs), a classic model for understanding principles of catalysis and enzyme design (*21-23*). These enzymes occur in bacteria, archaea, fungi, and plants and operate at the central branch-point of the shikimate pathway leading to the biosynthesis of tyrosine and phenylalanine (Fig. 1D). CMs catalyze the conversion of the intermediary metabolite chorismate to prephenate through a Claisen rearrangement, displaying more than a million-fold rate acceleration of this reaction (*24*), and are necessary for growth of bacteria in minimal media. For example, *Escherichia coli* strains lacking a CM are auxotrophic for Tyr and Phe, with both the degree of supplementation of these amino acids and the expression level of a reintroduced CM gene quantitatively determining the growth rate (*22*). Structurally, AroQα subfamily members exemplified by EcCM, the CM domain of the CM-prephenate dehydratase from *E. coli*, form a domain-swapped dimer of relatively small protomers (∼100 amino acids, Fig. 1E) (*25, 26*). Their size, essentiality for bacterial growth, and the existence of good biochemical assays makes AroQ CMs an excellent design target for testing the power of statistical models inferred from MSAs.

We used DCA to make a statistical model (Eq. 1) for an alignment of 1,259 natural AroQ protein domains that broadly encompasses the diversity of bacterial, archaeal, fungal, and plant lineages. Deducing the exact parameters (*h*_*i*_, *J*_*ij*_) from the observed statistics in the MSA (*f*_*i*_, *f*_*ij*_) for any protein is computationally intractable, but a number of approximation algorithms are available (*1*). Here, we use bmDCA, a computationally quite expensive but highly accurate method based on Boltzmann machine learning (*27*). For example, sequences generated by MC sampling from the model reproduce the empirical first- and second-order statistics of natural sequences used for fitting (Figs. 2A-B). More importantly, we also find that the model recapitulates higher order statistical features in the MSA that were never used in inferring the model (see Methods). This includes three-way residue correlations (Fig. 2C) and the inhomogeneously clustered phylogenetic organization of the protein family in sequence space (Fig. 2D). Our findings suggest that the statistical model goes beyond just being a good fit to the first- and second-order statistics to capture the essential rules governing the divergence of natural CM sequences through evolution. In contrast, a simpler profile model that retains only the intrinsic propensities of amino acids at sites (*h*_*i*_(*a*), fitted to reproduce frequencies of amino acids 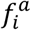) but leaves out pairwise couplings, fails to reproduce even the second-order statistics of the MSA (Fig. S1B), and does not account well for the pattern of sequence divergence in the natural CM proteins (Fig. S1D).

**Figure 2:**
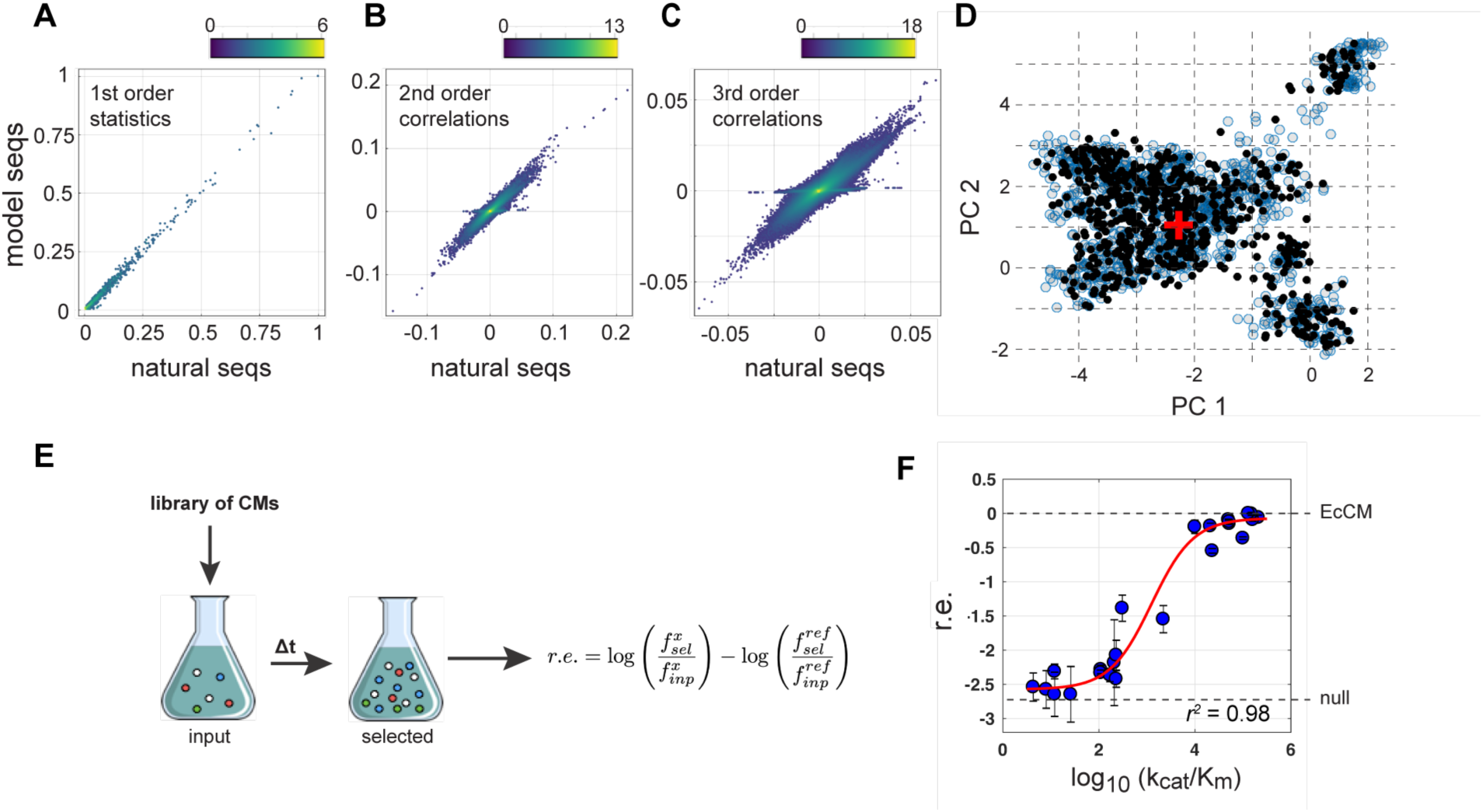
Design and testing of synthetic CM sequences. **A-C**, 2D histograms showing the relationship of first (A), second (B), and third (C) order statistics of natural and bmDCA-designed sequences. The color scale indicates the number of counts per bin. **D**, the top two principal components of the pairwise sequence distance matrix of natural homologs (blue circles) overlaid with a projection of synthetic CM sequences (black circles); the position of EcCM from *E. coli* is marked with a red plus sign. Synthetic sequences both recapitulate data used for fitting (A-B) and also account for statistical features of natural data not used for fitting (C-D). **E**, the workflow for functional characterization of chorismate mutase activity. CM deficient *E*.*coli* cells carrying libraries of variants are grown under selective conditions in minimal media, followed by deep sequencing of input and selected populations and calculation of the relative enrichment of each variant (r.e.). **F**, the relationship of r.e. to catalytic power (log_10_(k_cat_/K_m_)) for a number of CM variants yields a “standard curve”. The assay shows a hyperbolic relationship over the range from complete lack of CM activity to wild-type EcCM activity.

These findings raise the possibility that bmDCA may be a *generative model*, meaning that natural sequences and sequences sampled from the probability distribution *P*(***a***) are, despite considerable divergence, equivalent. To test this, we developed a high-throughput, quantitative *in vivo* complementation assay to monitor CM activity in *E. coli* that is suitable for studying large numbers of natural and designed CMs in a single experiment. Briefly, libraries of CM variants (natural and/or synthetic, see below) were made using a custom *de novo* gene synthesis protocol that is capable of fast and relatively inexpensive assembly of novel DNA sequences at large scale ((*28*) and see Methods). For example, we made gene libraries comprising nearly every natural CM homolog in the MSA (1,130 out of 1,259 total), and more than 1,900 synthetic variants exploring various design parameters of the bmDCA model (see Methods). These libraries were expressed in a CM-deficient bacterial host strain (KA12/pKIMP-UAUC, (*22*)) and all transformants grown together as a single population in selective media lacking phenylalanine and tyrosine to select for those variants exhibiting chorismate mutase activity (Fig. 2E). Deep sequencing of the population before and after selection allows us to compute the log frequency of each allele relative to wild-type EcCM. This quantity is called the “relative enrichment” (r.e.) which under particular conditions of gene induction, growth time, and temperature, quantitatively and reproducibly reports the catalytic CM activity (Fig. 2F, and Fig. S2). This “select-seq” assay is monotonic over a broad range of catalytic power and serves as an effective tool to rigorously compare large numbers of natural and synthetic variants for functional activity *in vivo*, in a single internally controlled experiment.

The first study examined the performance of the natural CM homologs in the select-seq assay as a positive control for bmDCA designed sequences. Natural sequences show a unimodal distribution of bmDCA statistical energies centered close to the value of EcCM (defined as zero, Fig. 3A), but it is not obvious how they will function in the particular *E. coli* host strain and experimental conditions used in our assay. For example, members of the CM family may vary in unknown ways with regard to activity in any particular environment, and the MSA includes some fraction of paralogous enzymes that carry out a related but distinct chemical reaction (*29, 30*). The select-seq assay showed that the 1,130 natural CM homologs exhibit a bimodal distribution of complementation in the assay, with one mode comprising ∼ 38% of the sequences centered close to the level of wild-type EcCM, and the remainder comprising a mode centered close to the level of the null allele (Fig. 3B). A Green Fluorescent Protein (GFP) tagged version of the library suggests that the bimodality of complementation is not obviously related to differences in expression levels compared to the *E. coli* variant (Fig. S3); instead, the bimodality presumably originates from non-linearities linking sequence to growth rate in the host strain, and from the inclusion of some functionally-distinct paralogous sequences. For the purpose of this study, the bimodality allows normalization of r.e. scores by the two modes and by Gaussian mixture modeling, to meaningfully group sequences into those that are functionally either like wild-type EcCM (norm. *r. e*. > 0.42) or like the null allele in our assay (Fig. 3B). Importantly, the standard curve shows that this quantity is a stringent test of high chorismate mutase activity (Fig. 2F).

**Figure 3:**
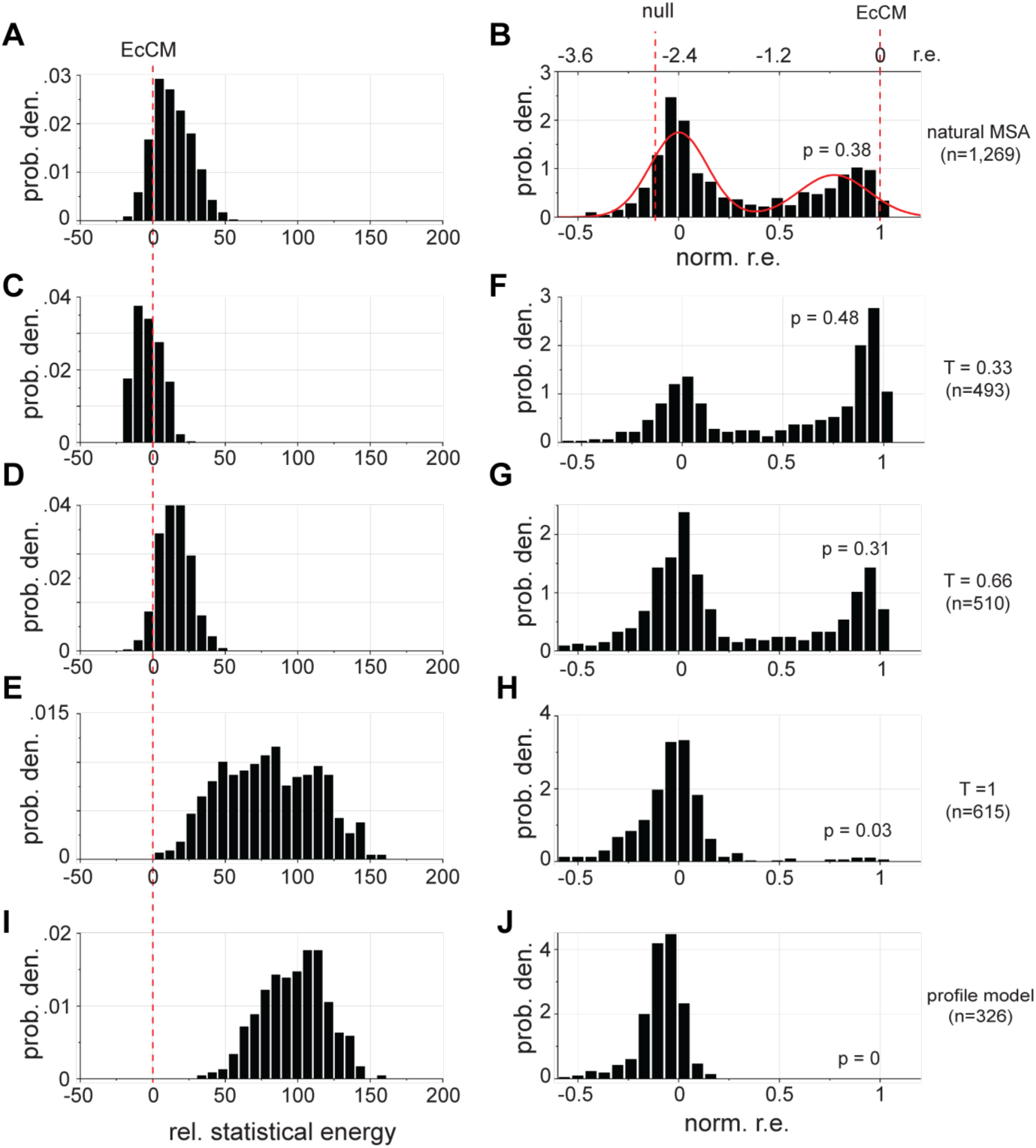
Functional analysis of natural and synthetic CMs. **A**, the distribution of bmDCA statistical energies for 1,130 tested natural AroQ homologs, relative to the value for EcCM. The data show a unimodal distribution centered close to EcCM. **B**, the distribution of functional complementation by natural AroQ sequences is bimodal, with ∼38% of sequences in one mode near to EcCM and the rest in another mode close to the r.e. of the null allele. The bimodality is used to normalize the raw r.e. scores between zero and the mean of the near-null mode for all libraries in panels B, F-H, and J. **C-E**, distributions of statistical energies for synthetic sequences and **F-H**, distributions of corresponding r.e. sampled at *T* ∈ {0.33,0.66,1}, respectively. **I-J**, distributions of statistical energy and r.e. for synthetic sequences retaining the intrinsic propensities of amino acids at positions but leaving out all correlations. Taken together, the data show that bmDCA is a generative model.

To evaluate the generative potential of the bmDCA model, we used MC sampling to randomly draw sequences from the model that span a wide range of statistical energies relative to the natural MSA (Figs. 3C-E), with the hypothesis that sequences with low energy (i.e. high probability) may be functional chorismate mutases. To sample sequences with low energy, we introduce a formal computational “temperature” *T* ≤ 1 in our model:

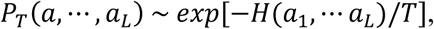

which, in exact analogy to temperature in statistical physics, serves to decrease the mean energy when set to values below unity. For example, sampling at *T* ∈ {0.33, 0.66} produces sequences with statistical energies that closely reflect the natural distribution (Fig. 3D) or reach even lower values (Fig. 3C). In contrast, sequences sampled at *T* = 1 show a broad distribution of statistical energies that deviate significantly from the natural distribution (Fig. 3E) towards higher energies. This deviation is, among other factors, due to statistical adjustments (regularization, see Methods) used during model inference for compensating for the limited sampling of sequences in the input MSA.

We made and tested libraries of 493-616 synthetic sequences sampled at *T* ∈ {0.33, 0.66, 1.0} from bmDCA models inferred at two regularization strengths (Fig. 3F-H). The data show that overall, these sequences also display a bimodal distribution of complementation, with many complementing function near to the level of the wild-type EcCM sequence. Consistent with our hypothesis, the probability of complementation is well-predicted by the bmDCA statistical energy, with low-energy sequences drawn from *T* ∈ {0.33, 0.66} essentially recapitulating or even somewhat exceeding the performance of natural sequences (Fig. 3F-G). In contrast, sequences drawn from *T* = 1 show poor performance, consistent with deviation in bmDCA statistical energies (Fig. 3H). Overall, 481 synthetic sequences out of 1,618 total (∼30%, norm *r. e*. > 0.42) rescue growth in our assay, comprising a range of top-hit identities to any natural chorismate mutase of 42 – 92% (Table S1 and Fig. S4A-B). These include 46 sequences with less than 65% identity to proteins in the MSA, corresponding to at least 33 mutations away from the closest natural counterpart. Sequence divergence from EcCM ranges from 14 to 82% (Fig. S4C-D). A representation of the positions in the EcCM protein that contribute most to the bmDCA statistical energy highlights residues distributed both within the active site and extending through the AroQ tertiary structure to include the dimer interface (blue spheres, Fig. 4F).

**Figure 4:**
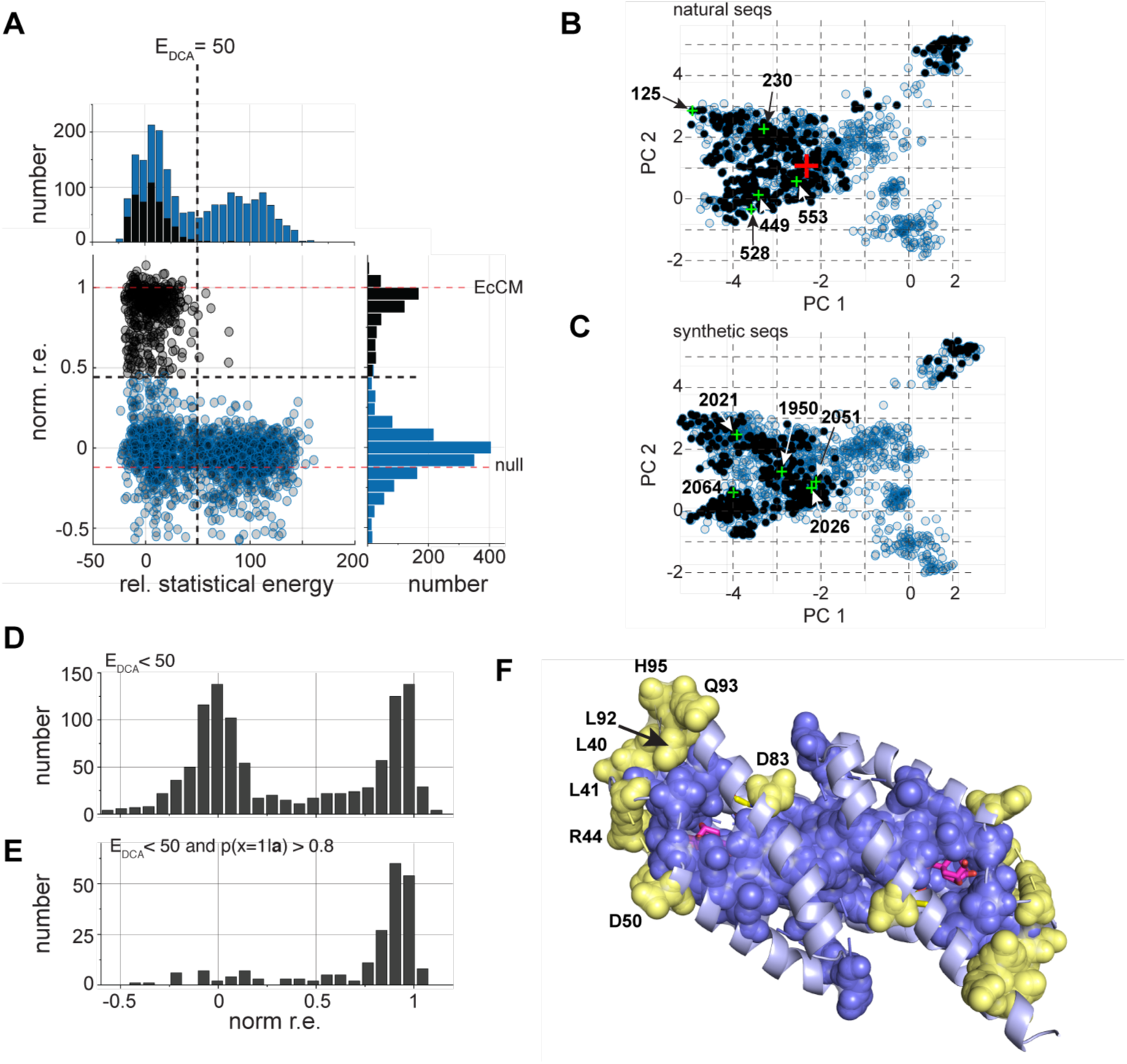
General and system-specific constraints in CM. **A**, the overall relationship of bmDCA statistical energies and catalytic function, with functional sequences in black and non-functional in blue. The data expose a steeply nonlinear relationship, with functional sequences strictly below a sharp threshold (*E*_DCA_ < 50). This value is the limit of statistical energies for natural homologs (Fig. 3A). **B**, the two top principal components of sequence variation in natural homologs, with sequences complementing the *E. coli* CM auxotroph in black. **C**, the same as panel b, but for synthetic sequences, showing a similar pattern. Sequences chosen for in-depth biochemical characterization are indicated in panels b-c, see Table 1. **D-E**, r.e. distributions for all synthetic sequences with *E*_DCA_ < 50 (d) or with an additional statistical constraint derived from the pattern of rescue of natural homologs (*P*(*x* = 1|***a***); see text for details). The additional constraint now identifies sequences functional in *E. coli* in our selection assay. **F**, The spatial architecture of functional constraints in CM enzymes mapped on to the EcCM structure. Blue spheres show positions constrained in the bmDCA model. Yellow spheres show the extra constraints required for *E. coli* specific function, highlighting a peripheral shell around active site residues important for CM catalysis.

Is the ability of designed CM sequences to rescue just a function of their sequence distance from their natural counterparts? To test this, we made 326 sequences with the same distribution of top sequence identities as bmDCA-designed sequences but preserving only the first-order statistics (position-specific conservation) and leaving out correlations. These sequences expectedly show high bmDCA energies and display no complementation at all (Fig. 3I-J). Thus, enzyme function is not simply about the magnitude of sequence variation and not even about conservation of sites taken independently; instead, it fundamentally depends on the pattern of correlations imposed by the couplings in *J*_*ij*_ (see Eq. 1).

**Table 1:** Biochemical properties of natural and designed CM enzymes. Construct numbers correspond to the numbering presented in Tables S1-S3, which provide additional information about these CM proteins.

| | Construct | Closest Natural | ID to EcCM | Top ID | $k_{cat}$ [$s^{-1}$] | $K_m$ [$\mu M$] | $k_{cat}/K_m$ [$M^{-1}s^{-1}$] |
| --- | --- | --- | --- | --- | --- | --- | --- |
| NATURAL | 1 | *Escherichia coli* | 1.00 | 1.00 | 64 | 390 | $1.6 \times 10^5$ |
| | | *Methanococcus jannaschii* | 0.30 | 1.00 | 5.7 | 41 | $1.4 \times 10^5$ |
| | 125 | *Nitratiruptor sp. SB155-2* | 0.32 | 1.00 | $110 \pm 10$ | $600 \pm 120$ | $2.0 \pm 0.6 \times 10^5$ |
| | 230 | *Geobacter uraniireducens* | 0.3 | 1.00 | $45 \pm 4$ | $37 \pm 7$ | $1.2 \pm 0.1 \times 10^6$ |
| | 449 | *Acetoanaerobium sticklandii* | 0.18 | 1.00 | $19 \pm 1$ | $190 \pm 30$ | $1.0 \pm 0.2 \times 10^5$ |
| | 528 | *Ilyobacter polytropus* | 0.24 | 1.00 | $19 \pm 2$ | $38 \pm 5$ | $5.1 \pm 1.2 \times 10^5$ |
| | 553 | *Collinsella intestinalis* | 0.23 | 1.00 | $7.6 \pm 3.2$ | $110 \pm 20$ | $6.8 \pm 1.8 \times 10^4$ |
| DESIGNED | 1950 | *Sulfurisphaera tokodaii* | 0.25 | 0.67 | $8.7 \pm 1.3$ | $120 \pm 10$ | $7.7 \pm 2.1 \times 10^4$ |
| | 2021 | *Desulfovibrio sp.* | 0.27 | 0.68 | $2.8 \pm 0.1$ | $330 \pm 10$ | $8.5 \pm 0.1 \times 10^3$ |
| | 2026 | *Methanocella arvoryzae* | 0.25 | 0.69 | $30 \pm 10$ | $220 \pm 30$ | $1.4 \pm 0.6 \times 10^5$ |
| | 2051 | *Methanococcus vannielii* | 0.22 | 0.66 | $3.0 \pm 0.1$ | $230 \pm 3$ | $1.3 \pm 0.0 \times 10^4$ |
| | 2064 | *Streptococcus sanguinis* | 0.22 | 0.69 | $21 \pm 13$ | $75 \pm 25$ | $2.7 \pm 0.8 \times 10^5$ |

We selected five natural and five designed CMs that complement growth in our assay for in-depth biochemical studies. The natural sequences occur in organisms representing a broad range of phylogenetic groups and diverse environments, and the designed sequences are chosen to sample regions that reflect this diversity (Fig. 4B-C). The selected CM genes were expressed in an *E. coli* strain that is deficient for endogenous CM to eliminate any possibility of contamination with the wild-type enzyme, and catalytic parameters of the purified proteins were determined using a spectrophotometric assay following the consumption of the substrate chorismate (*22*). The data show that all ten CM genes express similarly, and that the natural CMs display catalytic parameters similar to those of the previously characterized *E. coli(31)* and *M. jannaschii* (*32*) enzymes (Table 1). Consistent with their natural-like complementation, the designed CMs show catalytic parameters that closely recapitulate those of natural CMs. Thus, we conclude that the bmDCA-designed CMs are bona fide synthetic orthologs of the CM family.

Putting all the data together, we find a strikingly steep relationship between bmDCA statistical energy and CM activity (Fig. 4A and Fig. S5). Forty-five percent of designed sequences rescue the CM null phenotype when the statistical energy is below a threshold value set by the distribution of statistical energies observed for natural sequences (*E*_DCA_ < 50, Fig. 4A) and essentially no sequences (< 3%) are functional above this value. Thus, bmDCA infers an effective generative model, capable of designing natural-like enzymatic activity with considerable sequence diversity if statistical energies are within the range of natural homologs. The extent of sequence variation from natural homologs highlights the sparsity of the essential constraints on folding and biochemical function.

The bmDCA model captures the overall statistics of a protein family and does not focus on specific functional activities of individual members of the family. Thus, just like natural CM homologs, most bmDCA-designed sequences do not complement function under the specific conditions of our assay (Fig. 4A). But, might it be possible to improve the generative model to deduce the extra information that makes a protein sequence optimal for a specific phenotype? A bit of insight comes from studying how sequences that rescue function in our assay occupy the sequence space spanned by natural CM sequences. Natural CMs that complement function in our *E. coli* host strain are distributed in several diverse clusters (Fig. 4B), but interestingly, functional synthetic sequences also follow the same pattern (Fig. 4C). This suggests that information about CM function in the specific context of the *E. coli* assay conditions exists in the statistics of natural sequences and might be learned. If so, knowledge gained in the first experimental trial might be added to formally train a classifier to predict synthetic sequences that encode particular protein phenotypes and organismal environments.

To test this idea, we annotated the sequences in the natural MSA with a binary value *x* indicating their ability to function in our assay (*x* = 1 if functional, *x* = 0 if not). Due to its formal similarity to the DCA framework, we used logistic regression on the annotated MSA to learn a model providing a probability for any synthetic sequence ***a*** = (*a*_1_, ⋯, *a*_*L*_) to function in the *E. coli* select-seq assay; that is, *P*(*x* = 1|***a***). Interestingly, Figs. 4D-E show that for low-energy CM-like synthetic sequences sampled from the naïve, unsupervised bmDCA model (Fig. 4D), the extra condition that *P*(*x* = 1|***a***) > 0.8 now efficiently predicts the subset that complements in the context of our assay (83 %, Fig. 4E). These results support an iterative design strategy for specific protein phenotypes in which the bmDCA model is updated with each round of selection to optimize desired phenotypes.

What structural principles underlie the general constraints on CM function and the extra constraints for system-specific function? Mapping the positions that contribute most significantly to *E. coli*-specific function of CM sequences shows an arrangement of amino acids peripheral to the active site, within a poorly-conserved secondary shell around active site positions (Fig. 4F). Thus, these positions work allosterically or otherwise indirectly to control catalytic activity, a mechanism to provide context-dependent fine-tuning of reaction parameters.

The results described here validate and extend the concept that pairwise amino acid correlations in practically-available sequence alignments of protein families suffice to specify protein folding and function (*19, 20*). The bmDCA model is one approach to capture these correlations, but there is more work to be done to fully understand these models. Currently, the interpretation of DCA is focused on the relatively few highest magnitude terms in the matrix of couplings (*J*_*ij*_), because these identify direct structural contacts between amino acids in protein tertiary structures (*33*). Indeed, the top terms in *J*_*ij*_ for chorismate mutases do nicely correspond to contacts in the tertiary structure (Fig. S6). However, contact terms in *J*_*ij*_ alone do not suffice to reproduce either the alignment statistics in the AroQ family (Fig. S7) or the functional effects of mutations (*17*). Instead function in proteins seems to depend also on many weaker, non-contacting terms in *J*_*ij*_ that currently have no simple physical interpretation. Similar findings have been made in the case of predicting protein-protein interaction specificity (*34*). The weaker terms in *J*_*ij*_ seem to describe the collective evolution of amino acids within the structure, a property that may be related to patterns elucidated by other approaches to sequence coevolution (*1, 35-39*). A key next goal is to further refine the topology and best representation of sequence correlations that underlie the physics of protein structure and function.

However, even pending these necessary refinements, the data presented here provide the foundations for a general data-driven approach to protein engineering. This approach is similar to directed evolution in that it works without the use of physics-based potentials or atomic structures, but the computational models access a sequence space of functional proteins that is vastly larger than currently understood. Indeed, the results presented here permit a lower-bound estimate of the size of the sequence space consistent with the evolutionary rules for specifying members of a protein family. For example, at *T* = 0.66, we conservatively compute a total space of 10^25^ sequences that could be synthetic homologs of the AroQ family (see Methods). Given that ∼30% of sequences randomly sampled from this pool rescue CM function under our assay conditions, this amounts to more that 10^24^ sequences that can operate in a specific genomic and experimental context. These numbers are enormous in absolute terms but are infinitesimally unlikely in a sequence space searched without any model (∼10^125^) or with models capturing only first-order constraints (10^85^, see Methods). These considerations suggest that it will be of great interest to use evolution-based statistical models to guide the search for functional proteins with altered or even novel chemical activities.

## Acknowledgments

We thank O. Rivoire, C. Nizak, A. Ferguson, C. Feinauer, G. Uguzzoni, A. Pagnani, and members of the Ranganathan lab for discussions, A. George for software development, and the high-performance computing (BioHPC) and flow cytometry facilities at UT Southwestern Medical Center.

## Author contributions

W.R., M.W., R.M., S.C., and R.R. conceived the idea for the project, M.F., P. B-C., S.C., and M.W. built the computational models, W.R., with contributions from P.K. and D.H. developed the *in vivo* CM assay, W.R. built the synthetic gene libraries and carried out the *in vivo* assays, C.S., with contributions from M.S., P.K., and D.H. obtained the kinetic parameters in Table 1, S.C. and R. M. carried out the sequence entropy calculations, and W.R., M.W., and R.R. wrote the paper with contributions from all authors.

## Data and materials availability

Codes for bmDCA are available at https://github.com/ranganathanlab/bmDCA.

## Supplementary Materials

### Materials and Methods

Figures S1-S7

Tables S1-S3

