## Supplementary Information for "Evolution-based design of chorismate mutase enzymes"

.

**This PDF file includes:**

Materials and Methods
Figs. S1 to S7
Tables S1-S3
References

### Materials and Methods

Multiple sequence alignment:

Sequences were acquired by three rounds of PSI-BLAST (*40*) using residues 1-95 of EcCM (the CM domain of the *E. coli* CM-prephenate dehydratase, the P-protein) as the starting query (e-score cutoff 0.0001). We made an initial position-specific amino acid profile from a 3D alignment of four CM atomic structures (PDB entries 1ECM, 2D8E, 3NVT, 1YBZ) and then iteratively aligned nearest neighbor sequences from the PSI-BLAST search using the MUSCLE algorithm (*41*), each time updating the alignment profile and aligning nearest neighbor sequences from the PSI-Blast results. The resulting alignment was subject to a final round of hand adjustment using standard rules and then trimmed sequentially to retain positions present in *E. coli* CM (PDB ID 1ECM), to remove sequences with less than 82 residues, to remove sequences with more 30% gaps, and to minimize redundancy by removing excess sequences with more than 90% identity. The final MSA contains 1,259 sequences and 96 positions.

bmDCA model inference and sequence design:

The MSA was used to infer a Potts model, assigning a probability $P(a_1, \cdots, a_L) = \frac{1}{Z}\exp\{-H(a_1, \cdots, a_L)/T\}$ to each aligned sequence $(a_1, \cdots, a_L)$ with $L = 96$ (amino acids or gaps). The statistical energy (or Hamiltonian) $H(a_1, \cdots, a_L) = -\sum_{1\le i<j\le L} J_{ij}(a_i, a_j) - \sum_{1\le i\le L} h_i(a_i)$ of the Potts model is given in terms of the direct coevolutionary couplings $J_{ij}(a, b)$ between amino acids $(a, b)$ at positions $(i, j)$, and propensities (or fields) $h_i(a)$ for the usage of amino acid $a$ at position $i$. These parameters are inferred using bmDCA with reweighting threshold of 0.8 and regularization strengths $\lambda = 0.01$ or 0.001 (*27*). The formal temperature $T$ is set to unity during inference. The objective function in model inference is to accurately reproduce the empirical frequencies $f_i^a$ of each amino acid $a$ at each position $i$ and the joint frequencies $f_{i.j}^{ab}$ of amino acids $(a, b)$ at positions $(i, j)$, thus bringing positional conservation and covariation together:

$$f_i^{a_i} = \sum_{\{a_k|k\neq i\}} P(a_1, \cdots, a_L),$$

$$f_{ij}^{a_i a_j} = \sum_{\{a_k|k\neq i,j\}} P(a_1, \cdots, a_L).$$

To check the accuracy of the inferred model, we compare the sequence statistics of natural sequences with synthetic sequences drawn from $P(a_1, \cdots, a_L)$ by a Monte Carlo (MC) procedure. Specifically, we compare the connected two-residue correlations $C_{ij}^{ab} = f_{ij}^{ab} - f_i^a f_j^b$ and the three-residue correlations $C_{ijk}^{abc} = f_{ijk}^{abc} - f_{ij}^{ab} f_k^c - f_{ik}^{ac} f_j^b - f_{jk}^{bc} f_i^a + 2 f_i^a f_j^b f_k^c$ which describe the part of the empirical two- and three-residue frequencies that are not simply explained by the lower-order statistical quantities. The quantities $f_i^a$ and $C_{ij}^{ab}$ from MC samples represent stringent tests of the accuracy of fitting of the bmDCA models. Since protein MSAs are characterized by discrete, categorical (i.e. non-Gaussian) variables, one expects correlations of all orders from Potts models that are limited to just pairwise coupling terms. In that regard, there is no *a priori* reason why the three-residue correlations in the empirical input data should be reproduced. Thus, the fact that these higher-order statistics in the MSA are reproduced is a strong test of the generative nature of the bmDCA model.

The limited sampling of natural sequences ($\mathcal{O}(10^3)$), requires *regularized* inference to avoid overfitting the bmDCA parameters $\{J, h\}$; here we use $L_2$-regularization in which a penalty $\lambda\left(\sum_{1\leq i<j\leq L}\sum_{ab} J_{ij}(a,b)^2 + \sum_{1\leq i\leq L}\sum_a h_i(a)^2\right)$ is added to the likelihood of the data. The penalty systematically decreases the parameter values in the bmDCA inference, and thereby suppresses large parameter values resulting from poorly sampled rare events. This modifies the consistency equations during fitting:

$$f_i^{a_i} = \sum_{\{a_k|k\neq i\}} P(a_1, \cdots, a_L) + \lambda h_i(a_i),$$

$$f_{ij}^{a_i a_j} = \sum_{\{a_k|k\neq i,j\}} P(a_1, \cdots, a_L) + \lambda J_{ij}(a_i, a_j)$$

and has the effect of shifting the energy scale of MC sampled sequences from that of natural sequences:

$$\langle H\rangle_{nat} = \langle H\rangle_{MC} - \lambda\left(\sum_{1\le i<j\le L}\sum_{ab} J_{ij}(a,b)^2 + \sum_{1\le i\le L}\sum_{a} h_i(a)^2\right)$$

Thus, in our model, natural sequences have systematically lower energies than sampled sequences ($\langle H\rangle_{nat} < \langle H\rangle_{MC}$), requiring sampling at lower temperatures ($T < 1$) to produce sequences that have compatible energies with natural sequences. In practice, for the model with $\lambda = 0.01$ we find a difference of the average energies between natural and MC sequences of 74.8, cf. Figures 3A and 3E, while the regularization term accounts for a difference of 42.5, i.e. about 57% of the energy difference. For design, we sampled sequences from both regularization strengths $\lambda = \{0.001, 0.01\}$ as indicated in Table S1. To check that the MC procedure is sampling the sequence space in a statistically independent manner, we carry out multiple independent MC trajectories while checking that the autocorrelation of sampling within trajectories decays to the level of correlations between samples from the independent trajectories. From a practical point of view, bmDCA model inference and sequence design can be computed on desktop PC systems for the typical size of most protein domains (< 300 amino acids). Codes for bmDCA are available at https://github.com/ranganathanlab/bmDCA.

Gene construction:

*E. coli* codon-optimized genes coding for all natural and synthetic AroQ proteins were constructed using a PCR overlap extension method from a mixed pool of oligonucleotide fragments synthesized on microarray chips (Agilent). Two 230-mer oligonucleotides corresponding to each gene were designed with unique flanking primer annealing sites for “gene-specific primers” (GSPs, (*42*)) and a BtsαI (Type IIS) restriction site to remove the flanking region after amplification. Overlaps were at least 16 bases long with 3’ G or C bases and a melting temperature of at least 59°C (*43*). PCR was performed in 384-well plates using Q5 polymerase (NEB) with 1x Q5 buffer, 0.2 mM dNTPs and 0.5 µM GSPs in 10 µl total volume, with 35 cycles of 10 s each denaturation, annealing and primer extension at 98 °C, 61 °C and 72 °C, respectively. To remove the GSP annealing sites and amplify the full-length genes, the first round products were diluted

500x into a PCR reaction containing 0.1 U/µl BtsαI with flanking primers 5'-AGCGATCTCGGTGACGATGG-3' and 5'-CATTAACGATGCAAGTCTCGTGG-3' and incubated at 55 °C for 60 minutes prior to amplification for 10 cycles at 61 °C and 35 cycles at 65 °C annealing temperatures. Amplified products for all genes were pooled, digested with NdeI and XhoI, ligated into correspondingly digested plasmid pKTCTET-0 (*44*), column purified (Zymo Research), and transformed into electrocompetent NEB 10-beta cells (NEB) to yield >1000x transformants per gene. The entire transformation was cultured in 500 ml LB medium containing 100 µg/ml sodium ampicillin (Amp) overnight after which plasmids were purified, diluted to 1 ng/µl to minimize the likelihood of multiple transformation, and transformed into the CM-deficient strain KA12 containing the auxiliary plasmid pKIMP/UAUC (*22*) to yield >1000x transformants per gene. The entire transformation mixture was cultured in 500 ml LB containing 100 µg/ml Amp and 30 µg/ml chloramphenicol (Cam) overnight, supplemented with 16% glycerol and frozen at -80 °C.

Chorismate mutase selection assay:

Glycerol stocks of KA12/pKIMP-UAUC carrying pKTCTET variants with the CM gene libraries were cultured overnight at 30 °C in LB medium containing 100 µg/ml Amp and 30 µg/ml Cam. The culture was diluted to an $OD_{600}$ of 0.045 in M9c minimal medium containing 100 µg/ml Amp, 30 µg/ml Cam, and 20 µg/ml each of L-phenylalanine (F) and L-tyrosine (Y) (M9cFY, non-selective conditions), grown at 30 °C to an $OD_{600}$ of ~0.2, and washed with M9c (no FY). A portion of the washed culture was inoculated into 2 ml LB medium containing100 µg/ml Amp, grown overnight at 37°C and harvested for plasmid purification to generate the pre-selected ("input") sample. For the *in vivo* selection step, another fraction of the washed culture was diluted to a calculated starting $OD_{600}$ of $10^{-4}$ into 500 ml M9c containing 100 µg/ml Amp. 30 µg/ml Cam and 3 ng/ml doxycycline (for library gene induction from the $P_{tet}$ promoter on pKTCTET) and grown at 30 °C for 24 hours. The final $OD_{600}$ was < 0.1. Fifty ml of the culture was harvested by centrifugation, resuspended in 2 ml LB medium containing 100 µg/ml Amp, grown overnight at 37 °C, and harvested for plasmid purification (the "selected" sample).

Plasmids purified from input and selected cultures were amplified using two rounds of PCR with KOD polymerase (EMD Millipore) to add adapters and indices for Illumina sequencing. In the first round the DNA was amplified using primers that add from 6 to 9 random bases (Ns) for initial focusing, as well as part of the i5 or i7 Illumina adapters. In the second round of PCR, the remaining adapter sequence and TruSeq indices were added. For both rounds of PCR, only 16 cycles and a high initial template concentration were used to minimize amplification-induced bias. The final products were gel purified (Zymo Research), quantified using Qubit (ThermoFisher) and sequenced in an Illumina MiSeq system with a paired-end 250 cycle kit. Paired-end reads were joined using FLASH, trimmed to the NdeI and XhoI cloning sites and translated. Only exact matches to the designed genes were counted. Relative enrichment values were calculated according to equation $r.e. = log\left(\frac{f_{sel}^{x}}{f_{inp}^{x}}\right) - log\left(\frac{f_{sel}^{ref}}{f_{inp}^{ref}}\right)$, where $f_{y}^{x}$ is the frequency of observing gene $x$ in sample $y$, and the reference sequence (*ref*) is the gene for EcCM. The standard curve relating $r.e.$ to catalytic power (Fig. 2F) was made using published data for a set of 24 variants of EcCM, comprising A32S, A32T, E52A, E52D, E532Q, K39A, K39N, K39Q, K39R, L7C, L7V, Q88A, Q88E, Q88K, R11A, R11K, R28A, R28K, V35C, V35I, V35M, V85F, V85I, and wild-type (*45-47*).

<u>Chorismate mutase enzyme assays:</u>

pKTCTET plasmids encoding selected C-terminally $His_6$-tagged versions of natural and designed CMs were transformed into *E. coli* strain KA12/pT7POLTS (*44*). 100 ml of LB medium containing 100 µg/ml Amp and 30 µg/ml Cam were inoculated 1:100 with overnight starter cultures, grown at 37°C and 220 rpm to an $OD_{600}$ of 0.4-0.6, induced with 2 µg/mL tetracycline and further incubated at 20°C with shaking (200 rpm) overnight. Cells were harvested by centrifugation (4,000 g for 20 min at 4°C) and resuspended in 50 mM sodium phosphate, 0.3 M NaCl, pH 8.0, lysed with 1 mg/ml lysozyme for 30 min on ice and sonication (7-10 rounds with 70% amplitude, 0.7 cycle, 30 s each on ice using an UP 200 s tip, Sonotrode S7, Dr. Hielscher GmbH), and centrifuged at 20,000 g for 20 min at 4°C. CM proteins were purified from the cleared lysate by Ni-NTA affinity chromatography (Qiagen), dialyzed into a storage buffer (20 mM potassium phosphate, pH 7.5), and stored at 4°C. Chorismate was produced according to a published protocol (*48*). *In vitro* kinetic assays were performed as previously described(*22*). The depletion of chorismate was monitored at 274 nm ($\varepsilon_{274}$ = 2630 $M^{-1}$ $cm^{-1}$; substrate range 20-120 µM) or 310 nm ($\varepsilon_{310}$ = 370 $M^{-1}$ $cm^{-1}$; substrate range 50-600 µM) in 50 mM potassium phosphate buffer, pH 7.5, and 0.1 mg/mL BSA at 30°C, using a Perkin Elmer Lambda 35 UV/VIS spectrophotometer. The initial velocities ($v_0$) were determined and fitted into the Michaelis-Menten equation [$v_0 = (k_{cat}[E][S])/(K_m + [S])$] with KaleidaGraph (Synergy Software, Reading, PA, USA), where [E] and [S] are enzyme and chorismate concentrations, respectively**.** The rate constant $k_{cat}$ was calculated per CM active site. All parameters were calculated from the kinetics of at least two independently prepared biological replicates.

Analysis of gene expression: The library of natural CM sequences was cloned into a variant of the pKTCTET plasmid with 3' in-frame gene coding for the mNeonGreen fluorescent protein (*49*), transformed into KA12 cells, and grown at 30 °C in M9cFY medium (see above) containing 100 µg/ml Amp, 30 µg/ml Cam to $OD_{600}$ = 0.03. The culture was then transferred to a home-built turbidostat, grown at 30 °C with the $OD_{600}$ clamped at 0.05 for 17h to establish steady-state conditions, and then induced with 10 ng/ml doxycycline for 24h. We collected 1.6 million cells by fluorescence-activated cell sorting on a BD FACSAria (UT Southwestern cytometry core), selecting cells in the top 16.3% of fluorescence in the FITC channel. Both unsorted (input) and sorted (selected) cells were grown overnight at 30 °C in LB medium containing 100 µg/ml Amp. Plasmid DNA was purified from both input and selected populations, the genes encoding the cloned CMs were amplified by PCR, and the amplicons were gel purified and sequenced as described above. The relative expression level of each allele $x$ was calculated as the log ratio of the frequency of observing $x$ in the selected population relative to the input population, normalized by the same ratio for the wildtype EcCM gene. Only CM variants with at least 30 counts in the input population were used for analysis.

*Context-specific prediction:* To predict low-energy sequences that complement in the specific conditions of the *E.coli*-based assay, we annotate all natural sequences $\boldsymbol{a} = (a_1, \dots, a_L)$ with a binary value $x$ (1 if functional and 0 otherwise), and learn a model

$$P(x|\boldsymbol{a}) \sim \exp\left\{g \cdot x + \sum_i K_i(a_i) \cdot x\right\}$$

by logistic regression. In the model, $g$ is a general bias and $K_i(a_i)$ couples functionality $x$ to the specific amino acid $a_i$. The joint probability $P(x, \boldsymbol{a})$ takes thus the form of an extended DCA model in which the binary functional variable can be understood as an extra "position" added to each sequence of $L$ amino acids. After training using the annotated natural sequences, the conditional probability $P(x = 1|\boldsymbol{a})$ can be used to evaluate the assay-specific functionality of each designed sequence in the naïve MC sample.

Sequence entropy calculations:

Among the $20^{96} \approx 10^{125}$ sequences of length $L = 96$, only a small fraction correspond to functional CM proteins. To estimate their numbers, we present below several estimates of increasing accuracy.

We first consider an independent-site model, in which sequences $A = \{a_1, \dots, a_{96}\}$ are generated according to the factorized distribution $P(A) = \prod_{i=1}^{L} f_i^{a_i}$, where $f_i^{a_i}$ is the frequency of amino acid $a_i$ on site $i$, computed from the multi-sequence alignment (MSA) of natural sequences. The Shannon entropy of this distribution is

$$S_{independent} = -\sum_{i=1}^{L} \sum_{a_i} f_i^{a_i} \ln f_i^{a_i} \approx 198,$$

which corresponds to $e^{198} \approx 10^{85}$ sequences. This calculation grossly overestimates the number of functional CM sequences, as the independent model do not take into account the constraints coming from pairwise correlations between sites, crucial for the proper design of proteins.

Next, we compute the entropy of the bmDCA model, which includes those constraints. Due to the presence of couplings between the sites, an exact calculation of the entropy is not possible, and we resort to the so-called Annealed Importance Sampling (AIS) approximation method described below. The outcome is

$$S_{DCA} \approx 125 \pm 0.5,$$

which amounts to about 1.3 per site. This result is comparable to the values found in Barton et al. (*50*) for other protein families, e.g. 1.22 for the WW domain and 1.28 for the trypsin inhibitor Kunitz domain. The resulting estimate for the number of sequences is $e^{125} \approx 10^{54}$. Most of these sequences have, however, large energies and are not functional.

As shown in Fig. 3G, sampling the DCA-Potts model at temperature T=0.66 allows us to design functional CM proteins with a good probability. Using AIS again, we estimate the entropy at this low temperature to be

$$S_{DCA}(T = 0.66) \approx 58 \pm 1$$

or, equivalently, to about 0.6 per site. The corresponding total number of sequences is reduced to $e^{58} \approx 10^{25}$. All these results are summarized in the following Table (Statistical errors are standard deviations on the average values of the entropies over five repetitions of the AIS procedure).

| | All | Independent | bmDCA (T=1) | bmDCA (T=0.66) |
|---|---|---|---|---|
| Entropy (natural log) | 288 | 198 | 125±0.5 | 58±1 |
| Number of sequences | $10^{125}$ | $10^{85}$ | $10^{54}$ | $10^{25}$ |

We provide a brief description of entropy estimation through Annealed Importance Sampling (AIS). The entropy $S$ of the inferred model is related to its free energy $F$ and its average energy $U$ through the thermodynamic identity $S = (U - F)/T$, where $T$ denotes the temperature. Computing the average energy $U$ through Monte Carlo (MC) sampling of the inferred models is straightforward. The estimation of the free energy $F$ is more involved, and was done through AIS. Importance sampling (IS) is a method that allows one to accurately calculate the difference between the free energies of two similar models, through intensive

MC sampling of the probability of either of the models. We define a scheme of $M$ DCA models $m=0,...,M$ with increasing coupling strengths $\frac{m}{M} J_{ij}(a,b)$, interpolating between the independent model ($m=0$, no coupling, only fields) to the full DCA model ($m=M$). The free energy of the $m=0$-independent model can be easily calculated, and we use IS to estimate the free energy of the $m=1$ model, and so on, propagating along the chain of models until we can estimate the free energy of the $m=M$-full DCA model. This propagation scheme, consisting in annealing the coupling strength, is known under the denomination of AIS (*51, 52*), and was successfully tested on synthetic data in Barton et al. (*50*).

In practice, to achieve an accurate sampling of the models along the scheme by MC, at each step $m \rightarrow m+1$, we have run 500 parallel MC chains of length 100, using 2000 thermalization steps and 100 steps between each sample. For the annealing scheme, we have used $M=10^5$, and averaged over 100 annealing runs. The average entropies and their standard deviation given in Table above are computed over 5 repetitions of such computation.

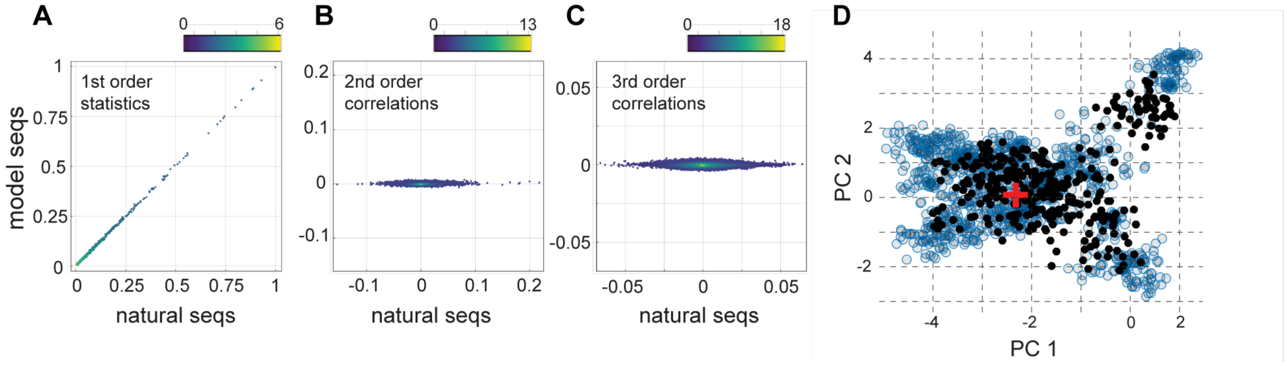


**Figure S1:** **A-C**, relationship of first (A), second (B), and third (C) order statistics of natural homologs to that from sequences sampled from a "profile model" with only first-order constraints (no couplings). **D**, the top two principal components of the pairwise sequence distance matrix of natural homologs (blue circles) overlaid with a projection of sequences drawn from the profile model (black circles); the position of EcCM is marked as a red plus sign. The data show that the profile model does not account for correlations in the natural MSA, and only partially recapitulates the pattern of sequence divergence of natural CM sequences.

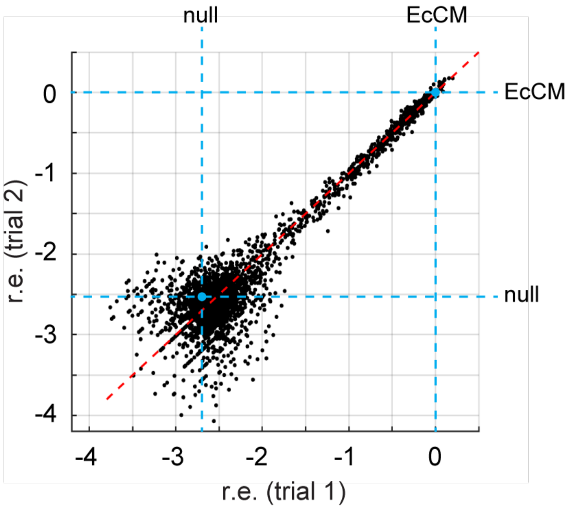


**Figure S2:** A scatterplot of relative enrichment (r.e.) values for two independent trials of the select-seq assay under the same experimental conditions for all 3074 CM sequences analyzed in this work (Tables S1-S3). The position of the wild-type EcCM sequence and the null allele (no CM activity) are indicated in blue circle and dashed lines as marked and the red dashed line is the identity trace. Values at low values of r.e. are subject to more variability as expected from poorer counting statistics. The data show that the select-seq assay shows excellent reproducibility.

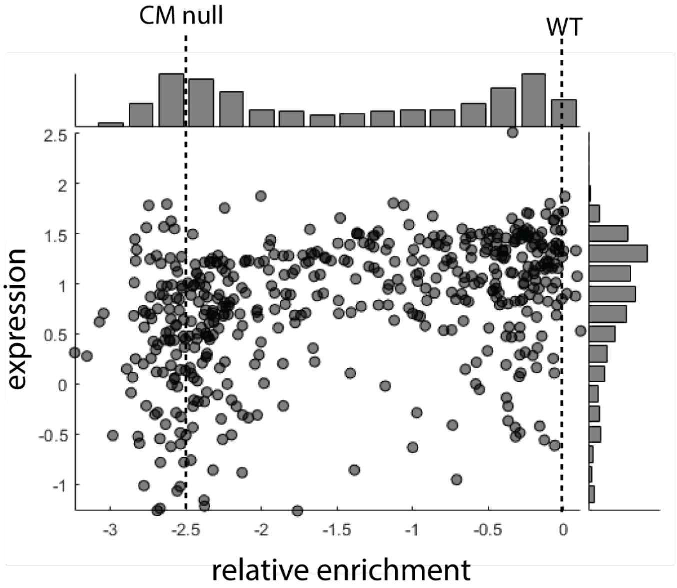


**Figure S3:** the relationship between functional rescue of natural CM enzymes and their expression levels in the *E. coli* host (CM null) strain. The expression level is followed by FACS analysis using a translational fusion of each natural CM with a fluorescent protein (mNeonGreen). The data show that the bimodality in functional complementation is not obviously explained by differences in expression.

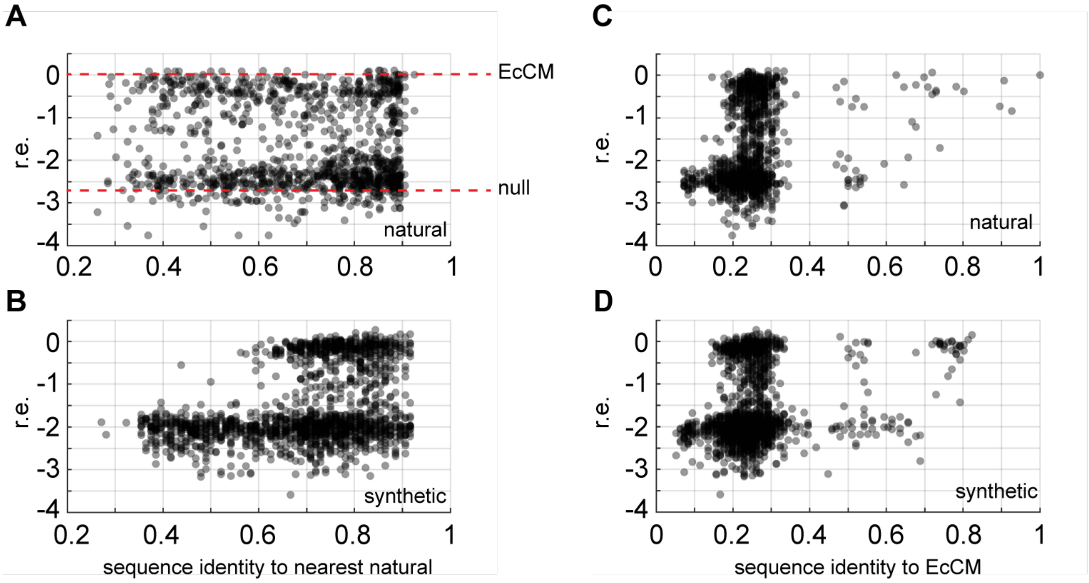


**Figure S4:** **A-B**, the relationship of relative enrichment (r.e.) to fractional sequence identity to the nearest natural sequence for sequences in the input MSA (A) and for all sequences sampled from the bmDCA model (B). **C-D**, the same plot but with relative enrichments plotted against fractional identity to the EcCM sequence.

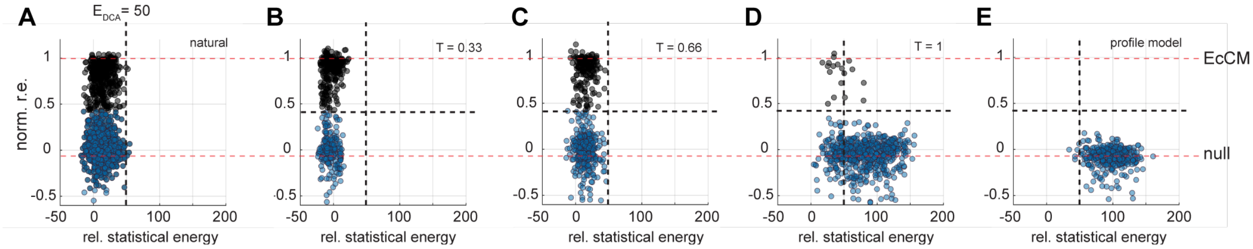


**Figure S5:** the relationship of relative enrichment to bmDCA statistical energy for the natural sequences (**A**), for sequences sampled from the bmDCA model at the three different computational temperatures (**B-D**), and for the profile model with no correlations (**E**).

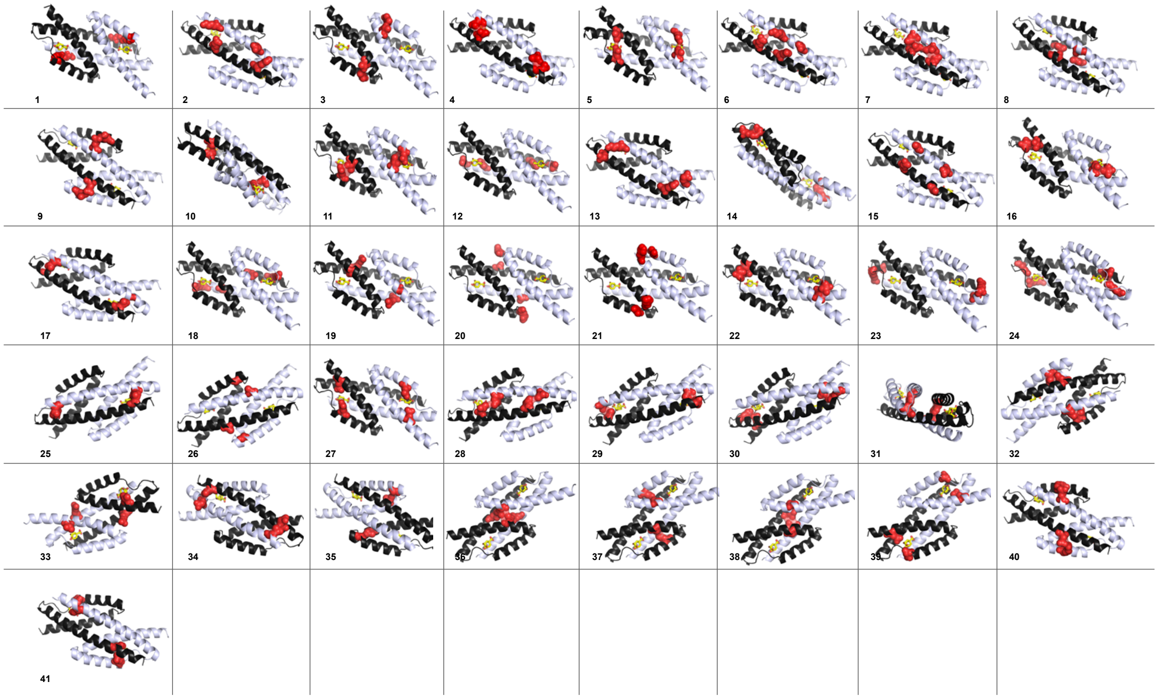


**Figure S6:** The top 41 pairwise amino acid interactions in the bmDCA model are shown as red spheres on the dimeric atomic structure of EcCM (PDB ID 1ECM(*25*)). The two CM protomers are shown in white or black cartoon representation. Nearly all statistically inferred couplings are direct contacts in the tertiary structure, either within one monomer or across the dimer interface. These data are consistent with the basic claims that the DCA model is an effective approach for contact prediction.

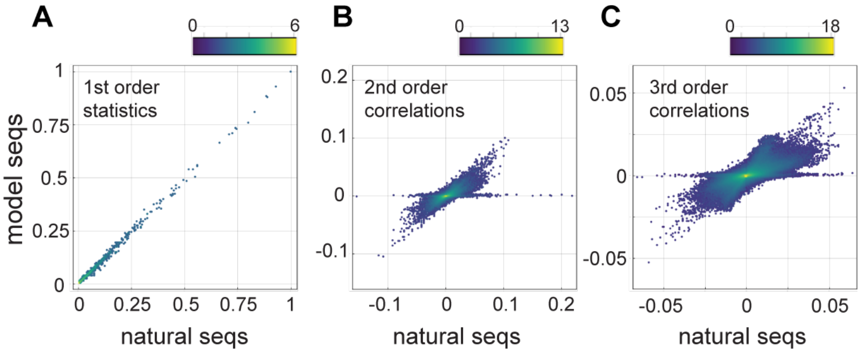


**Figure S7:** Relationship of first (A), second (B), and third (C) order statistics of natural homologs to that from sequences sampled from a bmDCA model keeping only direct contacts in the tertiary structure. The contacts-only model is poor at capturing the empirical statistics of those positions in the natural MSA, demonstrating the importance of the many non-contacting couplings for capturing the statistical features of natural sequences.

**Table S1:** The 1130 natural sequences comprising the input MSA. Each line contains the sequence, the species annotation, the DCA energy, the normalized relative enrichment, and the sequence identities to EcCM and to the closest natural sequence.

**Table S2:** The 1618 bmDCA sequences comprising those sampled at $T \in \{0.33, 0.66. 1\}$ and with models made with regularization strengths $\lambda \in \{0.01, 0.001\}$. Each line contains the sequence, the regularization value, the sampling temperature, the DCA energy, the normalized relative enrichment, and the sequence identities to EcCM and to the closest natural sequence.

**Tables S3:** The 326 sequences comprising the profile model (no correlations). Each line contains the sequence, the DCA energy, the normalized relative enrichment, and the sequence identities to *E. coli* and to the closest natural sequence.

Tables S1-S3 are provided as separated Excel files.
